# Rapid dissemination of host-metabolism-manipulating transposon-like entities via integrative and conjugative elements

**DOI:** 10.1101/2023.04.24.538000

**Authors:** Elena Colombi, Frederic Bertels, Guilhem Doulcier, Ellen McConnell, Tatyana Pichugina, Kee Hoon Sohn, Christina Straub, Honour McCann, Paul B. Rainey

## Abstract

Integrative and conjugative elements (ICEs) are self-transmissible mobile elements that transfer functional genetic units across broad phylogenetic distances. Accessory genes shuttled by ICEs can make significant contributions to bacterial fitness, yet ICEs that carry accessory genes encoding functions other than antimicrobial resistance remain poorly characterized.

Recent observation of the rapid acquisition of ICEs in a pandemic lineage of *Pseudomonas syringae* pv. *actinidae* led to investigation of the structural and functional diversity of these elements among a diverse array of *P. syringae*. Fifty-three unique ICE types were identified across multiple phylogroups. These ICEs display distinct evolutionary histories compared to their bacterial hosts, are highly recombinogenic, exhibit a conserved structure and are punctuated by hotspots of accessory gene integration. Many carry a 16 kb transposon-like entity (Tn*6212*) that shows little polymorphism indicating recent dissemination. Deletion of Tn*6212* did not alter pathogen growth *in planta*, but mutants displayed significant fitness defects when grown on TCA cycle intermediates. These were largely attributable to a single LysR regulator. RNA-seq analysis of a set of nested Tn*6212* deletions confirmed a central role of LysR in enhanced expression of more than 300 genes and down-regulation of genes controlling expression of energetically demanding loci. Together the transcriptional data indicate a major role for Tn*6212* in manipulation of bacterial metabolism with primary effects on RNA degradation, protein synthesis and potential diversion of ATP to growth.

## Introduction

Mobile genetic elements, such as plasmids and integrative and conjugative elements (ICEs), can move functional genetic units over broad phylogenetic distances, mediating abrupt changes in niche preferences and even contributing to speciation [1, 2]. Sequence analyses suggest that ICEs are the most abundant type of conjugative element in bacteria [3]. ICEs are plasmid-like entities with attributes of temperate phages that disseminate vertically as part of the bacterial chromosome, and horizontally by virtue of endogenously encoded machinery for conjugative transfer. During the process of conjugation ICEs excise and form circular intermediates. A conjugative relaxase introduces a single-strand nick and the ICE is then transferred to the recipient cell via a type IV conjugation apparatus. Site-specific integration of reconstituted double-stranded DNA occurs in the donor and recipient cell [4–6]. Genes encoding integration, excision, conjugation and regulation are typically encoded within modules referred to as ‘backbone’ loci [6]. In addition to essential genes, ICEs carry variable sets of accessory genes (or ‘cargo’) that make contributions to both ICE and host cell fitness. These include genes with functions associated with biofilm formation, pathogenicity and symbiosis, bacteriocin synthesis, antibiotic and heavy metal resistance [4, 5, 7].

*Pseudomonas syringae* is a model organism for the study of microbial evolution and plant-microbe interactions due to its ubiquity in both agricultural and non-agricultural areas. Different lineages of *P. syringae* are responsible for frequent outbreaks of disease in a variety of crop plants, and *P. syringae* can be found in association with wild plants, leaf litter, rivers, snowpack and even clouds [8–11]. *P. syringae* is more appropriately referred to as a species complex, comprising 13 divergent phylogroups (PGs) [12]. Although *P. syringae* is among the most well-studied bacterial plant pathogens, only eight ICEs have been described among the 901 complete and draft genomes of *P. syringae* species complex sequenced to date [13–16].

The recent emergence of a new lineage of *P. syringae* pv. *actinidae* (*Psa*) resulted in a global outbreak of bleeding canker disease on kiwifruit (*Actinidia* spp.), with severe consequences for agricultural production in Europe, Asia, New Zealand, Australia and Chile [17]. Population genomic analyses of *Psa* revealed the global outbreak was caused by a pandemic sublineage that emerged from a more diverse population of *Psa*-3 [18]. Separate introduction events of this clonal sublineage resulted in outbreaks in nearly all kiwifruit growing regions of the world. Initial genome comparisons showed the outbreak strains sampled from Italy, New Zealand and Chile, which varied by 6 single nucleotide polymorphisms (SNPs) in a 478,725 bp core genome, independently acquired three divergent ICEs during their global journey. The three ICEs have syntenic backbones sharing ∼75% nucleotide identity and carry identical 16 kb regions flanked by short palindromic sequences. Although lacking features typical of transposons, these mobile entities were labelled Tn*6212* [19] (and previously referred to as “enolase regions” [15]) and predicted to be linked to virulence of the pandemic sublineage of *Psa*-3 [15, 19]. After introduction of *Psa*-3 in New Zealand, where foliar copper sprays are frequently used to suppress infections, genomic surveillance revealed *Psa*-3 acquired a diverse pool of ICEs conferring copper resistance [16].

ICEs that carry accessory genes encoding functions other than antimicrobial resistance and nitrogen fixing symbiosis are relatively poorly characterized, even among mammalian pathosystems [6]. We first sought to determine the distribution and evolutionary history of ICEs in *P. syringae*, and to document the distribution of Tn*6212* . We identified a total of 207 ICEs present among six different phylogroups of the *P. syringae* species complex. This pool of ICEs comprises 53 distinct ICE types. Hotspots of cargo gene exchange were observed within otherwise conserved ICE backbones. Although a diverse cargo of accessory genes was identified, Tn*6212* was the most common cargo, present across 175 ICEs. We then sought to determine whether carriage of the transposon changes bacterial host phenotypes in plant-associated environments. We found Tn*6212* alters bacterial host gene expression, conferring a fitness benefit during growth on tricarboxylic acid cycle intermediates.

## Material and methods

### Identification and assembly of ICEs

A broad family of *P. syringae* ICEs (PsICEs) was identified using BLASTn searches of a collection of sequenced *Psa* genomes, combined with genomes deposited in the NCBI Genbank and WGS databases (updated to July 2021) [20]. When matches were identified in draft assemblies, contigs were downloaded and used to join contigs that overlapped by at least 6 bp. To delineate the chromosomal integration sites, flanking sequences were inspected for direct repeats (*att*) sites. The broader family of ICEs was defined with tBLASTn [20] searches in the NCBI GenBank WGS database (updated to April 2017).

### Identification and classification of PsICEs

REALPHY [21] was used to examine and classify PsICE diversity, identifying non-redundant PsICEs. REALPHY produced a final 1,975 bp alignment, which was in turn used to cluster similar PsICEs by building a UPGMA tree with 100 bootstrap replicates [22]. The UPGMA tree was used as a guide for the selection of a reduced set of representative, non-redundant PsICEs. The resulting set of 53 non-redundant PsICEs was used for subsequent analyses. The identification of conserved backbone genes in PsICEs was performed using the pangenome identification tool ROARY [23]. Backbone genes are here defined as genes present in at least 93% of all non-redundant PsICEs. MAFFT alignments using automatic alignment parameters [24] were used to examine structural conservation of the backbone genes and identify sites of accessory gene integration. Alignments were then separately generated for 62 backbone genes using MAFFT with automatic parameters [24]. Phylogenetic incongruence between individual backbone genes was evaluated using the ILD test [25] using PAUP* [26]. The extent of inter- PsICE recombination was evaluated with Neighbor-Net [27] using SplitsTree [28].

ClonalFrameML was also used on the concatenated alignment of the backbone genes [29]. Alfy 9 [30] was used to assess the inter-ICE recombination.

### Bacterial host phylogeny

A PhyML tree [31], using default parameters and 100 bootstrap replicates, was built using the concatenation of the alignments of the housekeeping genes *gapA*, *gltA*, *gyrB* and *rpoD* of each *P. syringae* genome, *P. fluorescens* SBW25 (AM181176.4) was used as outgroup.

### Deletion mutant generation, plant and bacterial growth conditions

*Pseudomonas* strains were routinely grown in KB at 28 ° C, *Escherichia coli* in LB at 37 ° C, and *Agrobacterium tumefaciens* in LB at 28 ° C. *Nicotiana benthamiana* and *Arabidopsis thaliana* Col0 assays were performed as previously described [32]. Deletion mutants in genes and regions of ICEPsaNZ13 (Table S7) were constructed by marker exchange mutagenesis as described in [33]. A *lacZ* reporter gene was introduced into *Psa* NZ13 for competition experiments via triparental mating as in [16].

### Bacterial protein secretion and host recognition assays

Two plasmids (pMT-1 and pMT-2) were constructed by Genscript® using vector pUCP22.

Each plasmid was introduced into both *Psa* NZ13 and *Psa* NZ13 Δ*hrcC* [33] via triparental mating. Binary vectors carrying the DctT:AvrRpt2 constructs were created to confirm truncated or full-length DctT did not interfere with AvrRpt2-mediated recognition of secreted proteins.

The *dctT:avrRpt2* was fused to a C-terminal epitope tag (3xFlag) and introduced into pICH86988 using Golden Gate cloning [34]. The plasmids were electroporated into *Agrobacterium tumefasciens* AGL1 as described in [32].

Hypersensitive response (HR) assays were carried out in *A. thaliana* Col-0 with strains diluted in 10mM MgCl_2_ to a final OD_600_ of 0.2 as described in [32]. The experiment was repeated twice. Ion leakage experiments were carried out as in [32]. *Agrobacterium* infiltration was used to transiently express DctT:AvrRpt2 constructs in *N. benthamiana*, along with *A. tumefasciens* strains carrying functional RIN4 and RPS2 expression constructs (AGLRIN4 and AGLRPS2, respectively) [35] as previously described [32].

### Competition and growth experiments

Competition experiments between wildtype *Psa* NZ13, *Psa* NZ13::*lacZ* and ICE mutant genotypes were performed *in vitro* using minimal M9 medium [36] supplemented with glucose (10 mM), fumarate (10 mM), citrate (10 mM), malate (20 mM), or succinate (10 mM) as sole carbon sources. 100 µl of cell washes at OD600 0.2 were used to inoculate 10 ml of M9 media (final OD600 of 0.002). Cultures were shaken at 250 rpm at 28°C, measuring bacterial density at 0, 2, 3 and 4 days by plating dilutions on KB amended with X-gal (60 µg ml^-1^) to distinguish between deletion mutants (white colonies) and ancestral *Psa* NZ13 marked with *lacZ* (blue colonies). The experiment was performed using three replicates and repeated three times. The fitness of each strain in the competition experiments is expressed as the Malthusian parameter [37].

### RNA extraction, sequencing and analysis

Strains were streaked to single colonies on KB plates and incubated at 28°C for 48-72 hours. Single colonies were used to inoculate 5mL of M9 medium supplemented with 20mM glucose, citrate or succinate. Liquid cultures were set up with 3 replicates per strain per carbon source, and shaken at 230 rpm at 28°C. Cultures were incubated for 26hrs (M9+succinate), 28hrs (M9+citrate) or 48hrs (M9+glucose). Cultures were then diluted into fresh media using three dilutions per sample to ensure collection at mid-log phase: 1:10, 1:25 and 1:50 (M9+glucose and M9+succinate) or 1:10, 1:20 and 1:40 (M9 + citrate). Cells were collected for RNA extraction at OD600 between 0.4 and 0.5. RNA was extracted using the RNeasy Mini Kit (Qiagen 74106) according to the manufacturer’s instructions. Samples were further treated with the Turbo DNA-free TM kit (ThermoFisher AM1907) according to manufacturer’s instructions. 210 ng total RNA was used for rRNA depletion with the bacterial Ribo-Zero kit

(Illumina 20037135), according to manufacturer’s recommendations. After rRNA depletion, the remaining RNA was fragmented, and Illumina-compatible libraries prepared using the NEBNext Ultra™ II Directional RNA Library Prep Kit for Illumina (New England Biolabs E7760S). Libraries were sequenced on the Illumina HiSeq3000 system. RNA sequencing data was analyzed as described in [38]. Genes with statistically significant differential expressions (p<0.05 and p<0.01) were annotated with KEGG pathways [39] and visualized using scripts stored at https://gitlab.gwdg.de/guilhem.doulcier/pseudomonas_rnaseq/. The results are available for browsing at http://www.normalesup.org/~doulcier/collab/2020_ICE/kegg/).

### Phenotypic characterization of motility on TCA cycle intermediates

Wild type and ΔTn*6212* mutant cells were grown in succinate and placed in Adler chambers with succinate or casamino acids (CAA) as attractants, and a PBS buffer control. Time-resolved data were collected by imaging at regular intervals (either 5 min intervals over the course of 1 h, or 500 ms over 15 s). The resulting images were subject to analyses that included measurement of cell swimming speed, directionality, and cell density at the moving swarm. The rate of radial expansion of both wildtype and ΔTn*6212* cells was measured by stab inoculating cells into semi-solid M9 agar containing either succinate (at pH 7.0 and pH 6.0), glucose (pH 7.0) or CAA (ph 7.0) as growth substrates. Stab inoculation was performed using initial cell density of either 10^5^ or 10^6^ cells, measuring radial expansion daily for 8 days using precision calipers.

The experiment was performed twice with, on each occasion, five replicates per treatment.

## Results

### An expanded family of ICEs is circulating in the *P. syringae* species complex

Bacterial whole genome sequences (updated to November 2017) and complete bacterial genomes (updated to July 2021) in NCBI GenBank were searched with BLASTn [20] using as query ICEPsaCL1, ICEPsaI10, ICEPsaNZ13, ICEPsyB728a and ICEPph1302A (Table S1) to identify homologous ICEs. This search resulted in a collection of 207 ICEs, collectively referred to as PsICEs (Table S1). The 207 PsICEs (Table S1) were found integrated adjacent to a tRNA-Lys gene, 41% in *att-2* and 32% in *att-1* (in the remaining 27%, the contig was too short to infer the position of the tRNA-Lys). The first attachment site (*att-1*) is proximal to *clpB* (*Psa* NZ13 IYO_024910), and the second site (*att-2*) is proximal to *queC* (*Psa* NZ13 IYO_008010) [15, 40]. ICEPs309-1 was integrated in a tRNA-Lys not adjacent to *clpB* or *qseC*. Although BLASTn searches were not restricted to any bacterial species, only ICEs present in plant-associated *Pseudomonas* spp. (i.e. *Pseudomonas syringae* species complex [11, 41]) were identified. PsICEs were harbored by diverse strains belonging to phylogroups (PG) 1, 2, 3, 4, 7 and 13 (Figure S1) [12]. PsICEs in PG1 strains are over-represented due to the availability of *Psa* genomes isolated from kiwifruit in China, South Korea and Japan [18] (McCann *et al*., *unpublished*). This constitutes a source of sampling-generated bias.

SNPs conserved across all 207 PsICEs were used to produce a cladogram (Figure 1).

**Fig. 1.**
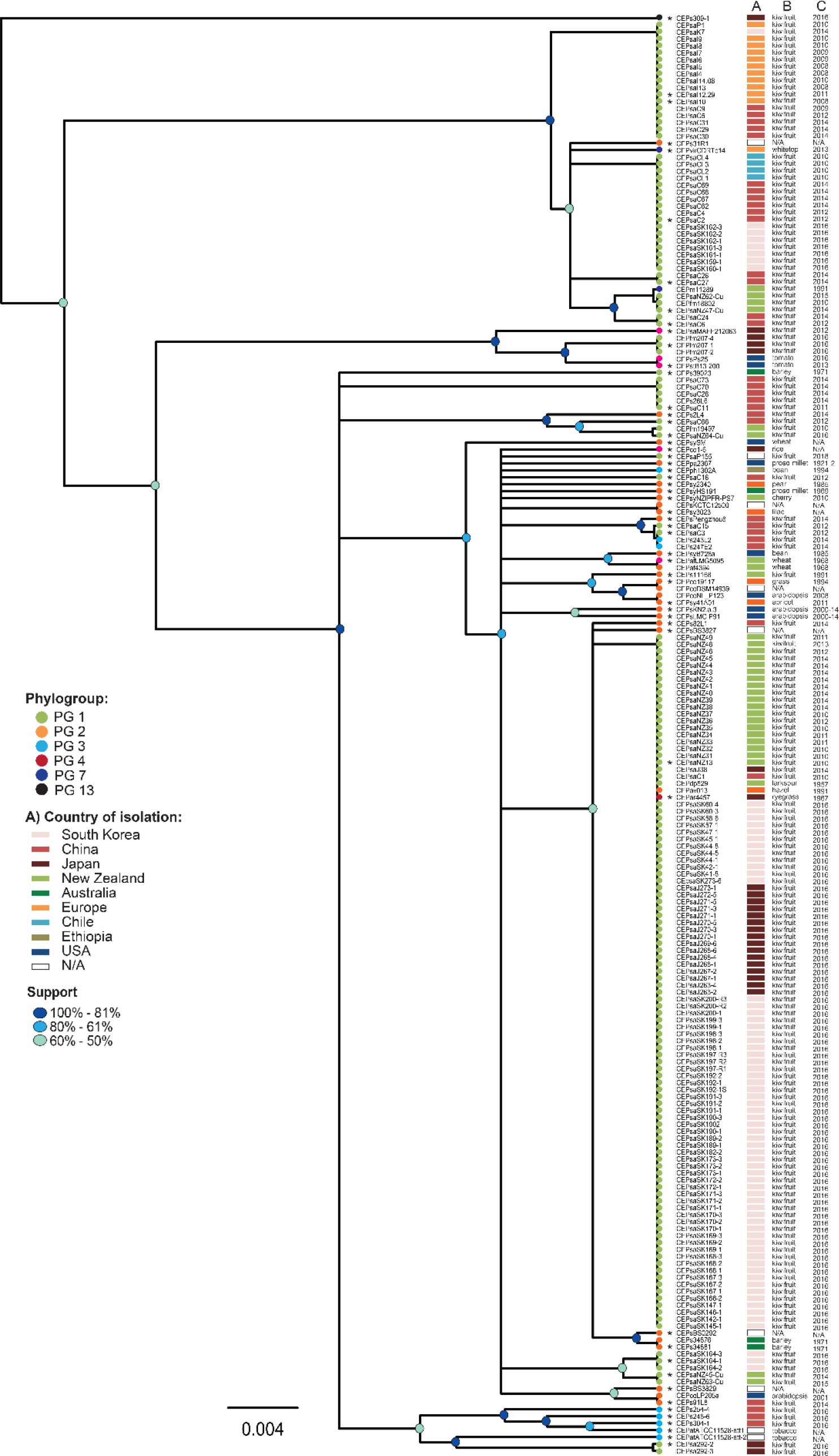
PsICEs are a large family of ICEs in the *P. syringae* species complex. An alignment of all PsICEs identified in the *P. syringae* species complex was generated using REALPHY. A 4.7kb alignment of conserved positions identified by REALPHY was then used to build a tree using UPGMA. Colors of terminal nodes indicate the phylogroup of the bacterial host genome harboring each PsICE, asterisks before PsICE names indicate non-redundant ICEs. Column A depicts the geographic location of the bacterial host, column B and C show the plant host from which the bacterium was isolated, and the year of isolation, respectively. Scale bar indicates substitutions per site.

While inter-ICE recombination [4, 7, 16] and the limited size of the aligned fragment (1,975 bp) mean that the resulting clusters should be treated with caution, it is notable that the largest cluster includes ICEs isolated from kiwifruit in New Zealand, China, Japan and South Korea from 2010 onwards. Interestingly, ICEs from divergent *P. syringae* isolated from larkspur (ICEPdp529) in 1957, hazel (ICEPav013) in 1991 and ryegrass (ICEPar4457) in 1967 fall into the same cluster. This cluster includes the canonical ICEPsaNZ13. ICEs within this cluster that share the same set of accessory genes as ICEPsaNZ13 (see below) are henceforward referred to as ICEPsaNZ13-like.

Consistent with capacity for horizontal transfer, PsICE distribution is incongruent with host strain phylogeny (Figure S1). Overall, PsICEs show no correlation with year, plant or geographic location of bacterial host (Figure 1). For example, ICEPsaNZ13-like elements are present in both divergent and pandemic sublineage *Psa*-3 strains: in PG1 (ICEPsaNZ13, ICEPsaC1), PG2 (*P. syringae* pv. *avellanae* ISPaVe013) and PG4 (*P. coronafaciens* pv. *atropurpurea* ICMP4457). ICEPsaC15-like and ICEPsaC3-like elements are highly similar, and have been isolated in both *P. syringae* PG1 (*Psa*) and *P. savastanoi* (PG3) host strains found in association with kiwifruit in China. Conversely, distinctly different ICEs are present in otherwise closely related host strains isolated from the same plant host, in the same geographical area.

ICEs from different years sometimes cluster together, for example, the aforementioned ICEPdp529 (1957) and ICEPar4457 (1967) group with ICEPsaNZ13-like elements from 2016.

To identify ICEs in *Pseudomonas* genomes other than those of *P. syringae*, the NCBI GenBank repository of *Pseudomonas* genomes (excluding *P. syringae*) was interrogated in April 2017 with the ICEPsaNZ13 DEAD-box helicase protein sequence, the most highly conserved gene among all PsICEs. Forty-four ICEs carrying DEAD-box helicases were identified in 41 *Pseudomonas* genomes, 82% of which were *P. aeruginosa* (Table S4). All ICEs were integrated in one of the two *att* sites described for PsICEs, with the exception of an ICE in *P. aeruginosa* PA38182, which did not harbor recognizable *att* sites. Thirty-three ICEs were part of the pKLC102/PAGI-2 family of ICEs [42] (Table S2). Although the pKLC102/PAGI-2 and PsICEs form clearly distinct families (Figure S2), conserved ICE life cycle genes are shared among the two families and are syntenic (Figure S2). This finding thus places the PsICEs in a broader context of ICEs found in gammaproteobacteria [43].

### PsICEs conserve their structure despite frequent inter-ICE recombination

ICE sequences are typically composed of conserved backbone genes, predicted to be involved in the ICE life cycle, and variable accessory (or cargo) genes. Identification of the PsICE backbone was guided by identification of the set of genes present among at least 93% of all ICEs using ROARY [23]. The backbone consists of a set of 62 genes (∼56 kb) predicted to be involved in ICE maintenance, regulation and movement, as well as a number of conserved hypothetical proteins (Table S3 and Figure S3). Homologs of these genes in other ICEs have been shown to encode conjugation machinery (*pil*), ICE transfer (*tra*), partitioning (*par*) and integration (*int, xerC*) functions [6]. The relaxed criterion chosen here (present in at least 93% ICEs) reflects the possibility of misassembly or gene deletion events (Figure S3). PsICE backbone genes are syntenic, with an average nucleotide identity across all genes of 86.4%. Average nucleotide identity varies between 94.1% for the gene encoding the DEAD-box helicase and 74.8% for a hypothetical protein-encoding gene (backbone gene #27). When distantly related PsICEs are compared (e.g., ICEPsaNZ13 vs ICEPsaI10 and ICEPsaNZ13 vs ICEPs309-1), the pairwise identity of the DEAD-box helicase gene decreases to 92.7% and to 70.3%, respectively (Figure S3).

Despite conservation of genes required for core ICE function and mobility, signatures of recombination are evident among ICEs. Backbone gene trees display phylogenetic incongruity (ILD tests, *P* of type I error = 0.01) [25]. ClonalFrameML [29] detects several recombination events in an alignment of concatenated backbone genes, and Neighbor-net [27] produces a highly reticulated network with a statistically significant Phi test for recombination (P<0.0001) (Figure S4) [44]. Finally, an alignment-free method of sequence comparison [30] indicates each PsICE is a chimera of other PsICEs, with short stretches of sequence displaying no homology to any known PsICE (Table S4). Thus, inter-ICE recombination appears to have shaped PsICE evolution and diversity, largely obscuring the evolutionary history of the PsICEs.

### Variable cargo genes are present in PsICE insertion hotspots

A subset of 53 non-redundant PsICEs was identified from the initial 207 PsICEs based on their position in the UPGMA tree and on differences in gene content within clusters (Table S1, Figure 1). This set of non-redundant PsICEs was used for all subsequent analyses. Comparison of the 53 non-redundant PsICEs reveals the backbone serves as a scaffold for variation introduced in ten specific positions, referred to as cargo regions (CR). The CRs are found in intergenic positions, with the exception of CR9, an integration hotspot likely driven by the presence of *rulAB* [45] (Figure 2). The integration of genes in CR9 results in the partial deletion of *rulA* or *rulB.* EGGNOG functional prediction [46] of the complete set of cargo genes shows 70% have no predicted function. Some cargo genes are notably abundant however: Tn*6212* is integrated into CR4 in 30 of the 53 non-redundant PsICEs. Imperfect direct repeats found at the extremities of Tn*6212* [19] may constitute sequences recognized by the XerC site-specific tyrosine recombinase. Clusters of heavy-metal resistance genes integrated in CR4, alone, or in tandem with Tn*6212*, are surrounded by the same repeats. Arsenic resistance genes are present in 13 non-redundant PsICEs; copper (and cadmium) resistance genes are found in 8, and two ICEs (ICEPaf4394 and Ps34881) harbor a ∼7 kb transposon encoding mercury resistance. In contrast, PsICEs harbor few type 3 secretion system (T3SS) effectors (T3SE): *hopAR1* is present in ICEPph1302A, ICEPs304-1 and ICEPfm207, ICEPatATCC11528-att2 carries *hopF2*, *hopO1-1*, *hopT1-1*, ICEPsaMAFF212036 harbours *hopAU1*, and ICEPs248-6 captured *avrRmp1* in the *rulAB* hotspot.

**Fig. 2.**
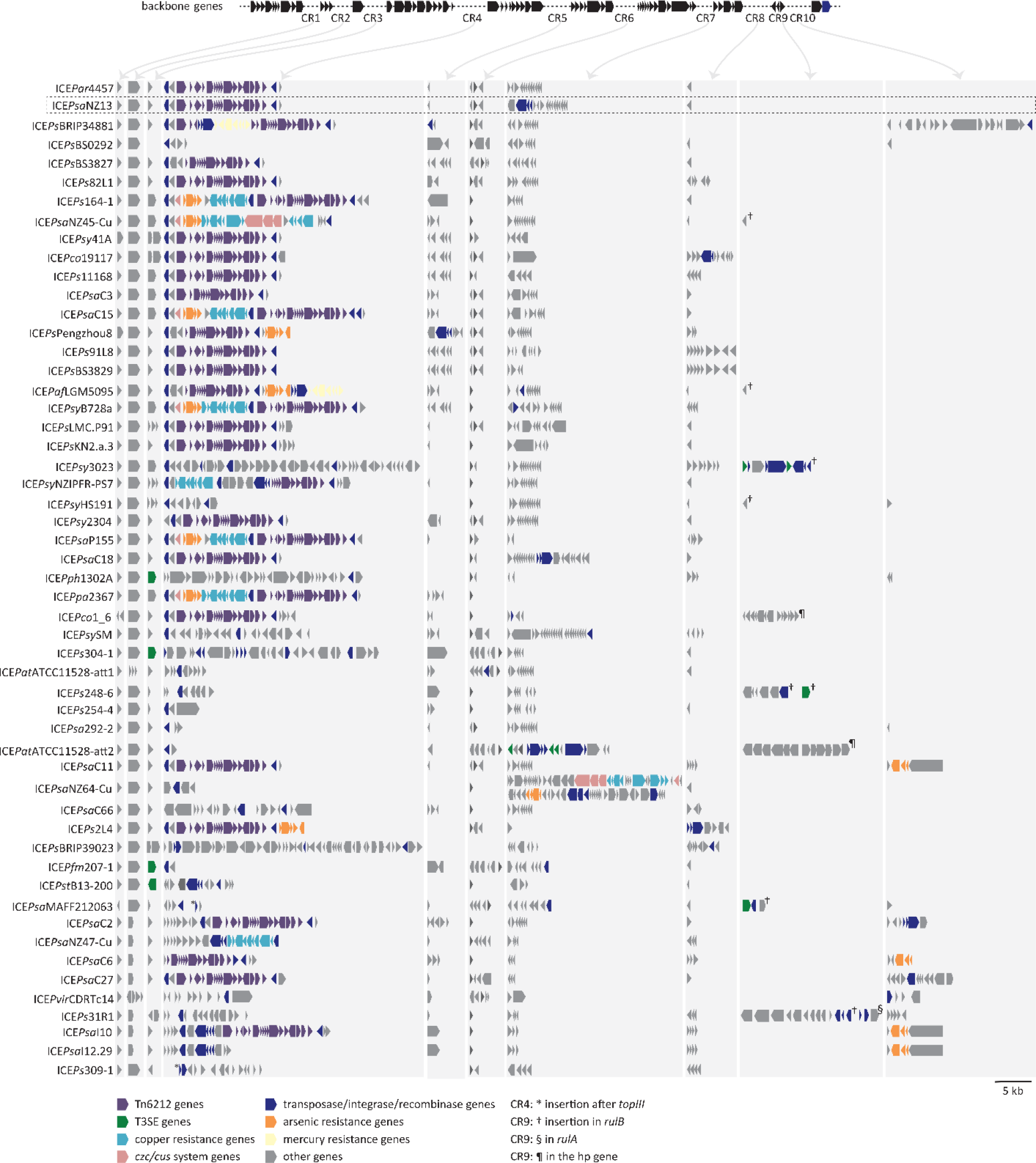
Genetic organization of PsICE backbone and hotspots of cargo gene integration. The gene content of the CR is highlighted with grey background; arrows indicate their position on the backbone. The distance between contiguous backbone genes varies accordingly to the content of the CR. ICE*Psa*NZ13 is highlighted with a rectangular box.

### Tn*6212* enhances the growth of *Psa* NZ13 on TCA cycle intermediates

The high frequency and nucleotide identity of Tn*6212* across otherwise divergent PsICEs suggest the transposon has recently spread and may confer a fitness advantage in plant-associated bacteria. Tn*6212* is a ∼16 kb tyrosine recombinase transposon that consists of a set of 20 genes (7 coding for hypothetical proteins) (Table S5). While the full length Tn*6212* is the most common cargo element, eight PsICEs carry only subsets of the Tn*6212* genes (Figure S5). Although PsICE backbones often share low levels of pairwise identity, the full length Tn*6212* transposons share over 99% pairwise nucleotide identity.

The genes encoded by Tn*6212* are not obviously associated with pathogen virulence or antibacterial resistance. However, Tn*6212* encodes genes implicated in the transport and metabolism of organic acids, including genes predicted to encode an enolase, a pyrophosphatase and a transporter of dicarboxylic acid (DctT). *P. syringae* is known to use reverse carbon catabolite repression and prefers organic acids as carbon sources [47]. The presence of a T3SS-targeting signal led to the hypothesis DctT might be exported via T3SS into plant cells to deprive the plant of C4 sugars [15]. To determine whether DctT is exported, the *dctT* promoter and its putative T3SS-targeting signal (1-52 aa) was fused to the C-terminal sequence of *avrRpt2* (pMT1), a T3SE recognized by *Arabidopsis thaliana* Col-0 [48]. A second construct (pMT2) included the promoter and full length *dctT*. Both *dctT:avrRpt2* constructs were confirmed as functional using transient expression via agroinfiltration into *Nicotiana benthamiana* (Figure S6). After introduction into *Psa* NZ13 and *Psa* NZ13Δ*hrcC,* which lacks a functional T3SS [33], strains were inoculated into *A. thaliana*, and plants were monitored for AvrRpt2 recognition via ion leakage assays or the development of a hypersensitive response (Figure S6). These experiments showed DctT was not exported via the T3SS, or via other means, in *Psa* NZ13. We then investigated whether carriage of Tn*6212* was involved in bacterial growth on plant hosts, but *Psa* NZ13ΔTn*6212* was not significantly impaired in growth compared to wildtype after flood inoculation of *A. chinensis* var. *chinensis* Hort16A (Figure S7).

Although there was no evidence of DctT secretion, and no *in planta* phenotype was detected, the presence of genes whose products are associated with energy production and sugar utilization (enolase *eno*, inorganic pyrophosphatase, a catabolism associated protein *cta*, and *dctT*) nevertheless suggested that Tn*6212* is associated with bacterial growth and metabolism. To test this hypothesis, competitive fitness assays between *Psa* NZ13 and *Psa* NZ13ΔTn*6212* were performed in M9 minimal media supplemented with glucose or TCA cycle intermediates (citrate, succinate, malate or fumarate) as sole carbon sources. *Psa* NZ13ΔTn*6212* showed a significant reduction in fitness compared to the wildtype in M9 containing TCA cycle intermediates, but not in glucose (Figure 3, Figure S8). In order to identify the genes responsible, three non-overlapping deletion mutants were generated within different regions of Tn*6212* (Figure 3): *Psa* NZ13 Tn*6212*Δ1 (∼6kb deletion including 8 genes), *Psa* NZ13 Tn*6212*Δ2 (∼6kb deletion including 7 genes), *Psa* NZ13 Tn*6212*Δ3 (∼4kb deletion including 3 genes and *xerC*) and *Psa* NZ13 Tn6*212*Δ*dctT* (Table S5). *Psa* NZ13 Tn*6212*Δ3 displayed a growth deficit comparable to *Psa* NZ13ΔTn*6212* in all TCA intermediates, so further single gene mutants in this region were created (*Psa* NZ13Δ*lysR*, Δ*cta*, Δ*lpr*). *Psa* NZ13Δ*lysR* and *Psa* NZ13Δ*cta* exhibited reduced growth on all TCA intermediates, although this reduction was not comparable to the growth deficit exhibited by the deletion of the entire transposon (Figure 3, Figure S8). Curiously, *Psa* NZ13Δ*lpr* exhibited enhanced growth in M9 supplemented with citrate and malate. It thus seems *lysR* and *cta* interact synergistically with effects that are distinct from those caused by Δ*lpr*.

**Fig. 3.**
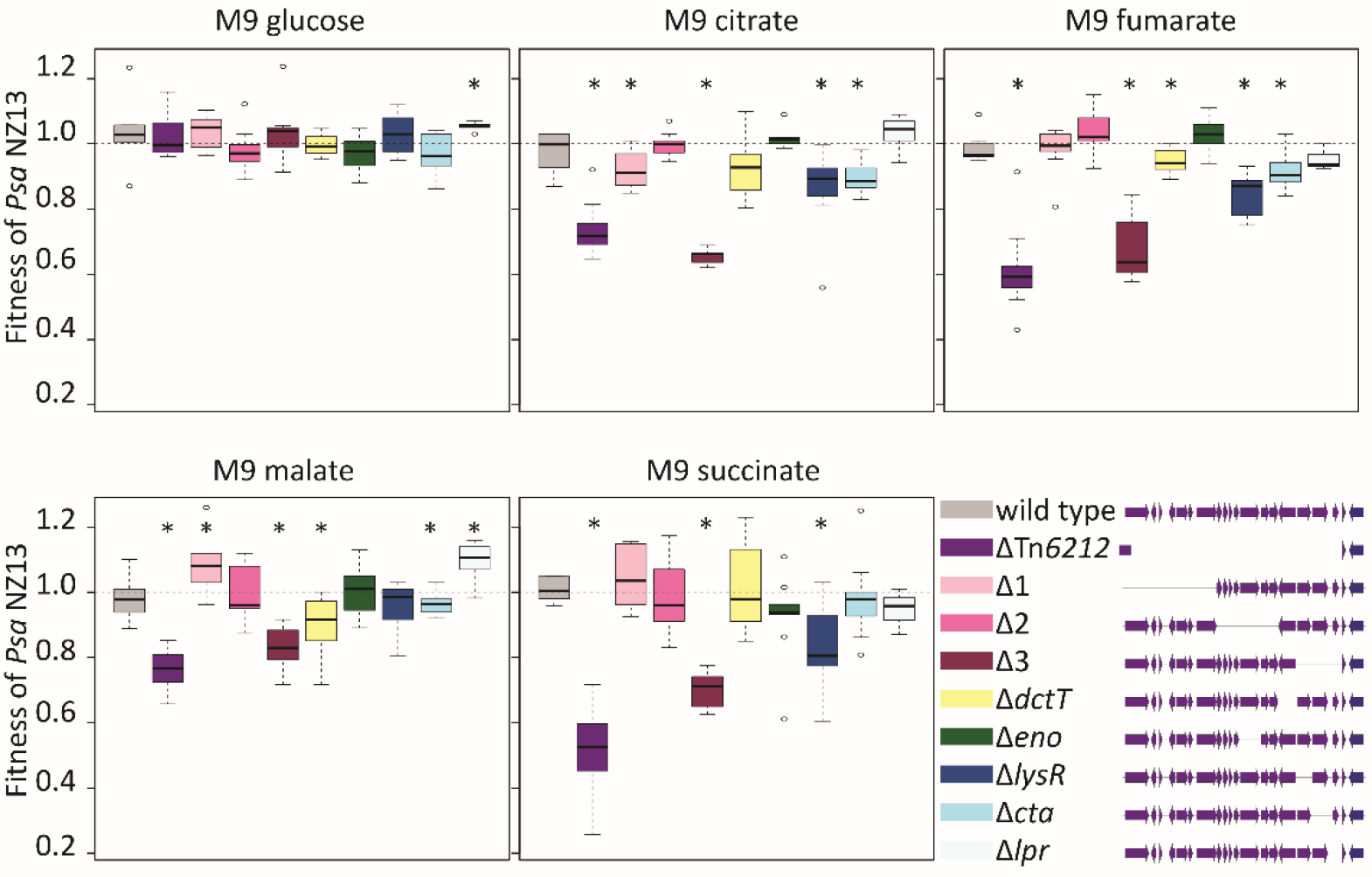
Competition assays. Box plots showing the fitness of Tn*6212* mutants relative to *Psa* NZ13 *Psa* NZ::tn7-*lacZ*. Competition assays (1:1) were performed in M9 supplemented with different carbon sources, only day 4 is shown. Data for day 2, 3 and 4 are shown in Figure S8. Values smaller than 1 indicate a lower relative fitness of competitor. The experiment was performed with three replicates and repeated three times. From left to right, *Psa* NZ13 wild type, *Psa* NZ13ΔTn*6212*, *Psa* NZ13 Tn*6212*Δ1, *Psa* NZ13 Tn*6212*Δ2, *Psa* NZ13 Tn*6212*Δ3, *Psa* NZ13Δ*dctT*, *Psa* NZ13Δeno, *Psa* NZ13Δ*lysR*, *Psa* NZ13Δ*cta*, *Psa* NZ13*Δlpr*. * indicates that the difference in fitness is statistically significant (one-sided one sample t-test p < 0.05).

### Tn*6212* has global effects on *Psa* NZ13 gene regulation

After observing the contribution of Tn*6212* to *Psa* NZ13 fitness on TCA cycle intermediates, we asked whether Tn*6212* has an impact on the regulation of PsICE activity, and whether Tn*6212* instigates broader changes consistent with manipulation of host cell metabolism. The latter seemed conceivable given the presence of the versatile and promiscuous LysR regulator [49]. RNA-seq was performed on *Psa* NZ13 and a set of nested deletion mutants (*Psa* NZ13ΔTn*6212*, *Psa* NZ13 Tn*6212*Δ3, *Psa* NZ13Δ*lysR*), plus *Psa* NZ13Δ*dctT*. All strains were grown in M9 with glucose, citrate and succinate as sole carbon sources, and RNA extractions performed once cells reached late exponential phase of phase of growth.

Transcriptional responses of each mutant were compared to wildtype *Psa* NZ13 grown in the same media and genes and transcriptional responses were annotated with KEGG [39].

A plausible null expectation is that strains cluster based on carbon source. The Euclidean distance plot displaying the normalized mean expression of the wildtype and mutants grown in M9 supplemented with glucose, citrate or succinate shows this is only the case when strains are grown in glucose (Figure 4A). *Psa* NZ13ΔTn*6212*, Tn*6212*Δ3 and Tn*6212*Δ*lysR* mutants share more similar expression profiles with each other when grown on citrate and succinate than with the wild type strain (Figure 4A). This indicates the absence of Tn*6212*; *lysR*, *cta* and *lpr*; and *lysR* alone results in significant differences in expression compared to wildtype during growth on citrate and succinate. There is greater similarity between Tn6212Δ*dctT* and wildtype expression on TCA intermediates than between Tn6212Δ*dctT* and other Tn*6212* mutants. This is at odds with the fact that the Δ*dctT* mutation is nested within Tn*6212* and thus expected to have just a subset of the effects wrought by the entire transposon. Its clustering with wild type, *Psa* NZ13, suggests that the transcriptional effects of *dctT* are distinct from those caused by the totality of genes on Tn*6212*.

**Fig. 4.**
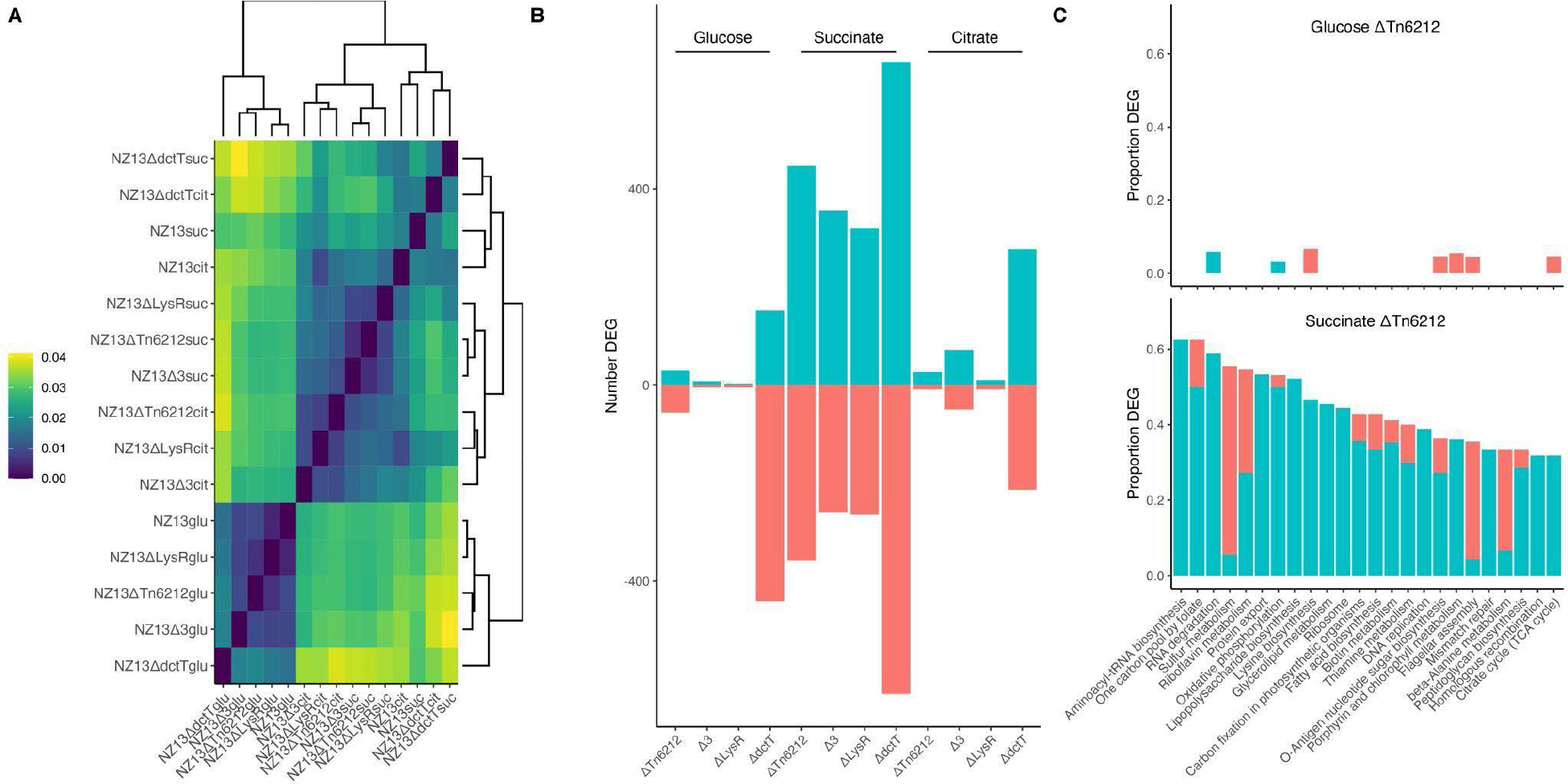
Tn*6212* alters bacterial host transcriptional responses (A) For each RNA seq dataset the normalized mean coverage of every open reading frame encoded in the chromosome was calculated using Deseq2 [50]. The Euclidean distance between the datasets was also calculated with each dataset being represented as a vector of normalized mean expression values. A distance of 0.04 means that expression differs on average by 4% per gene. The datasets cluster by carbon source used for growth except for NZ13 wildtype and NZ13Δ*dctT* grown in succinate and citrate, which cluster by genotype rather than carbon source. (B) Number of differentially expressed genes in each of the genotypes grown in different carbon sources. Negative values indicate the number of genes that are significantly (p ≤ 0.05) under expressed, positive values the number of genes that are significantly overexpressed compared to the NZ13 wildtype grown in the same carbon source. (C) KEGG functional categories that are significantly over- (turquoise) or under expressed (red) in NZ13ΔTn*6212* compared to the wildtype when grown in glucose and succinate. All functional categories containing at least 10 genes and where at least 30% of those genes are significantly differentially expressed are shown. We also show level 2 categories from the following top level KEGG hierarchies: Environmental Information Processing, Cellular Processes, Genetic Information Processing and Metabolism. For example, over 60% of all genes in the Aminoacyl-tRNA biosynthesis pathway are overexpressed in NZ13ΔTn*6212* when grown on succinate compared to the NZ13 wildtype. In contrast there is not a single gene in the same pathway that is differentially expressed when grown in glucose.

The number of differentially expressed genes (*P* < 0.05) and the relationship with genetic background and carbon source shows that the number of differentially expressed genes is greatest for strains grown on succinate (Figure 4B). The magnitude of effects extends well beyond both Tn*6212* and the ICE, indicating that Tn*6212* manipulates host cell metabolism.

Leaving aside *dctT*, whose deletion had no effects on fitness at later time points, the majority of regulatory effects can be directly attributed to *lysR*: in succinate, of the 792 genes whose expression is significantly altered on deletion of Tn*6212*, 411 are also affected in *Psa*NZ13Δ*lysR* (r = 0.978, p < 0.001) (Figure S9). For each carbon source, the number of differentially expressed genes is highest in *Psa* NZ13Δ*dctT*. This indicates that DctT functions, either directly or indirectly, as a repressor of genes on Tn*6212*. Given that DctT is a predicted transporter for di-carboxylic acids, it seems likely that information gathered via the transporter, perhaps concerning the nature of the external environment, is necessary for Tn*6212* to coordinate ensuing effects on host cell gene expression.

Making sense of the myriad transcriptional changes poses a major challenge. Of particular interest are those genes causally responsible for the observed changes. Figure 4C shows the proportion of genes differentially expressed by Tn*6212* when grown on succinate and connection to various cellular functions as defined by the KEGG database resource [39]. Data are ranked by the proportion of genes with significantly altered patterns of expression and the entire dataset with graphical mapping to KEGG pathways can be viewed at http://www.normalesup.org/~doulcier/collab/2020_ICE/kegg/. The number of genes whose expression is differentially affected by Tn*6212* when grown on glucose, for the same set of KEGG pathways, is shown for comparison. When grown on succinate, Tn*6212* significantly, and with primarily positive effects, affects the expression of numerous genes in multiple KEGG pathways involved in translation (tRNA biosynthesis, one carbon pool by folate, sulphur metabolism, protein export, ribosome), post transcriptional control (RNA degradation), energy metabolism (oxidative phosphorylation, sulfur metabolism), carbohydrate metabolism (amino sugar and nucleotide sugar biosynthesis, TCA cycle), metabolism of cofactors and vitamins (thiamine, riboflavin, and biotin metabolism) and DNA metabolism (DNA replication, mismatch repair, homologous recombination).

Repressive effects are few, with notable exceptions being in sulfur metabolism, flagella assembly (and chemotaxis) and beta-alanine metabolism. Closer inspection of the affected genes shows that for sulfur metabolism, the activity of genes involved in production of sulfate or sulfite are decreased, whereas genes contributing to the synthesis of homocysteine and thus methionine show enhanced expression. For beta-alanine metabolism, genes responsible for conversion of beta alanine to 3-oxopropanoate and then acetyl-CoA are repressed, but expression of pantoate-beta-alanine ligase, which converts beta alanine to D-4- phosphopantothenate (and thus pantothenate) is significantly increased. The major repressive effect is on flagella assembly and chemotaxis where numerous genes are significantly repressed, although the magnitude of change is low (∼1.5-fold decrease).

### Motility of Tn*6212* mutants on TCA cycle intermediates

We attempted to connect observed alterations in gene regulation to phenotypic changes by focusing on genes contributing to motility and chemotaxis. Measurement of cell swimming speed, directionality, and cell density in Adler chambers showed no significant differences between the mutant and wildtype genotypes. Having failed to detect differences in cell-level behavior, we then examined the rate of radial expansion of wildtype and ΔTn*6212* genotypes stab-inoculated with 10^5^ cells into semi-solid M9 agar containing either succinate (at pH 7.0 and pH 6.0), glucose (pH 7.0) or casamino acids (CAA) (pH 7.0) as growth substrates. No significant difference in the rate of radial expansion was detected on M9 glucose (*P*=0.97) or CAA (*P* = 0.25), but on M9 succinate, carriage of Tn*6212* significantly increased the rate of radial expansion, with the highest rate evident at pH 6.0 (*P* = 0.009 and *P* < 0.001, at pH 6.0 and pH 7.0, respectively). We observed substantial variability among all genotypes in growth initiation time using an initial density of 10^5^ cells on glucose. Whereas growth on succinate or CAA was readily detectable by 1 day for all replicates starting at 10^5^ cells, the initiation of growth in glucose was highly variable, requiring at least 4 days, and in one replicate no growth was visible even at the final time point of 8 days. Suspecting density-dependent behaviour, an additional treatment was included in which 10^6^ cells were used to found the centrally located population. This increased inoculum density largely eliminated variation in growth initiation and showed that the presence of the transposon decreased the rate of radial expansion on glucose (Figure S10).

## Discussion

ICEs associated with the *P. syringae* species complex were first identified in 2000 as “pathogenicity islands” whose spontaneous excision caused switches in virulence phenotypes of *P. syringae* pv. *phaseolicola* [51]. Since 2000, awareness of ICEs as vehicles that move ecologically significant genes has rapidly grown, with particular evidence of diversity and horizontal transfer coming from genomic analysis of *Psa* strains associated with the global kiwifruit canker disease pandemic [15, 19], and dramatic evidence of their impact arising from study of copper resistant strains in New Zealand [16]. Here, our bioinformatic analyses show that the *P. syringae* complex harbours numerous and diverse ICEs distributed across a range of phylotypes. Although core genes show overall synteny, genes of presumed ecological relevance are located in defined cargo regions. A large number of ICEs carry Tn*6212*, which contributes positive fitness effects on TCA sugars. Recent global dissemination appears connected to the capacity of the transposon to manipulate host cell metabolism.

Comparative analysis clearly shows ICEs are facilitators of horizontal transfer, mediating movement of diverse sets of genes among a wide range of bacterial hosts and over significant spatial scales. Near-identical PsICEs were found in hosts sampled decades and thousands of kilometers apart, and near identical host strains carry PsICEs with backbone genes that share less than 95% average nucleotide identity, a threshold commonly used to define distinct bacterial species [52]. While PsICEs are defined by a conserved set of core genes, all ICEs carry genes likely to confer ecologically significant functions to hosts. In some instances, these show matches to known genes, but many are either function-unknown, or have no orthologues in databases. Of note is the relatedness of PsICEs to those from *P. aeruginosa* raising the possibility that ICEs move genes between plant and opportunistic human pathogens, although minimally-overlapping ecological niches likely limit ICE movement across larger phylogenetic distances.

Many PsICEs carry accessory genes that confer a selective advantage to agricultural plant pathogens, like antimicrobial resistance and virulence-associated genes. The identification of cargo genes relevant to the bacterial host niche is not unusual, STX-R391 ICEs of clinical origin typically carry antibiotic resistance genes [53], while STX-R391 ICEs from free living marine bacteria *Alteromonas* mainly encode for metal resistance and restriction modification systems [54]. We have previously shown copper resistance genes on an ICE confer a growth advantage to *Psa* on leaf surfaces sprayed with copper [16]. The widespread application of bactericides, insecticides and herbicides is likely to select for the maintenance of copper and arsenic resistance genes after ICE acquisition. Type 3 secreted effectors (T3SE), known to play an important role in disrupting plant host recognition and immune responses, are also present on ICEs. The very first ICE characterized in the *P. syringae* species complex (ICEPph1302A) carries *hopAR1*, which is also present on ICEPs304-1 and ICEPfm207. ICEPs248-8 carries *avrRpm1*, ICEPsaMAFF212036 harbours *hopAU1*, and ICEPatATCC11528-att2 carries three effector genes: *hopF2*, *hopO1-1* and *hopT1-1*. Notably, all of these effectors are known to elicit effector-triggered immune responses. The transfer of a recognised T3SE onto an ICE may allow the pathogen to evade plant host recognition by silencing or maintaining the virulence gene at low frequencies in the population [55]. The most common cargo carried by PsICEs is not associated with agricultural sprays or plant host resistance responses, but is rather a transposon (Tn*6212*) associated with the transport, regulation and metabolism TCA cycle intermediates.

The widespread occurrence and high sequence identity of Tn*6212* from diverse ICEs indicates it recently invaded PsICEs and likely confers a selective advantage to *P. syringae*. We show here that Tn*6212* enhances bacterial fitness on TCA cycle intermediates, which are abundant in plant tissues and upregulated during pathogen invasion [56]. The Tn*6212* C4- transporter DctT and transcriptional regulator LysR may sense shifts in carbon source availability and initiate a signaling cascade altering the expression of chromosomal genes. The widespread impact on chromosomal expression is a challenge to interpret, however the examination of patterns of altered expression, functional categories and connections to genes carried on Tn*6212* provide some clues that the mechanism behind the altered fitness caused by Tn*6212* resides in RNA degradation.

Tn*6212* encodes the glycolytic enzyme enolase, a major component of the RNA degradosome connecting the physiological status of the cell to RNA degradation [57], and enolase is significantly over-expressed in the *Psa* NZ13 Tn*6212*Δ3 mutant (and with elevated expression in Δ*lysR* (*P* = 0.90)). Tn*6212* also encodes an inorganic pyrophosphatase that contributes to RNA degradosome function with the possibility of additional contributions from uracil-DNA glycosylase. Beyond genes encoded by the transposon, all major additional components of the RNA degradosome, including RNAse E and two different DEAD/DEAH box helicases are significantly over-expressed in *Psa* NZ13 growing on succinate, compared with ΔTn*6212*. Further reason to suggest a causal role of the RNA degradosome stems from its principal role in post transcriptional control. As a secondary consequence, the expression of genes involved in TCA cycle and gluconeogenesis is stimulated. Thus, it appears that Tn*6212* is able to detect preferred carbon sources and rapidly redistribute incoming carbon to maximize ATP, promoting growth (Figure 5).

**Fig. 5.**
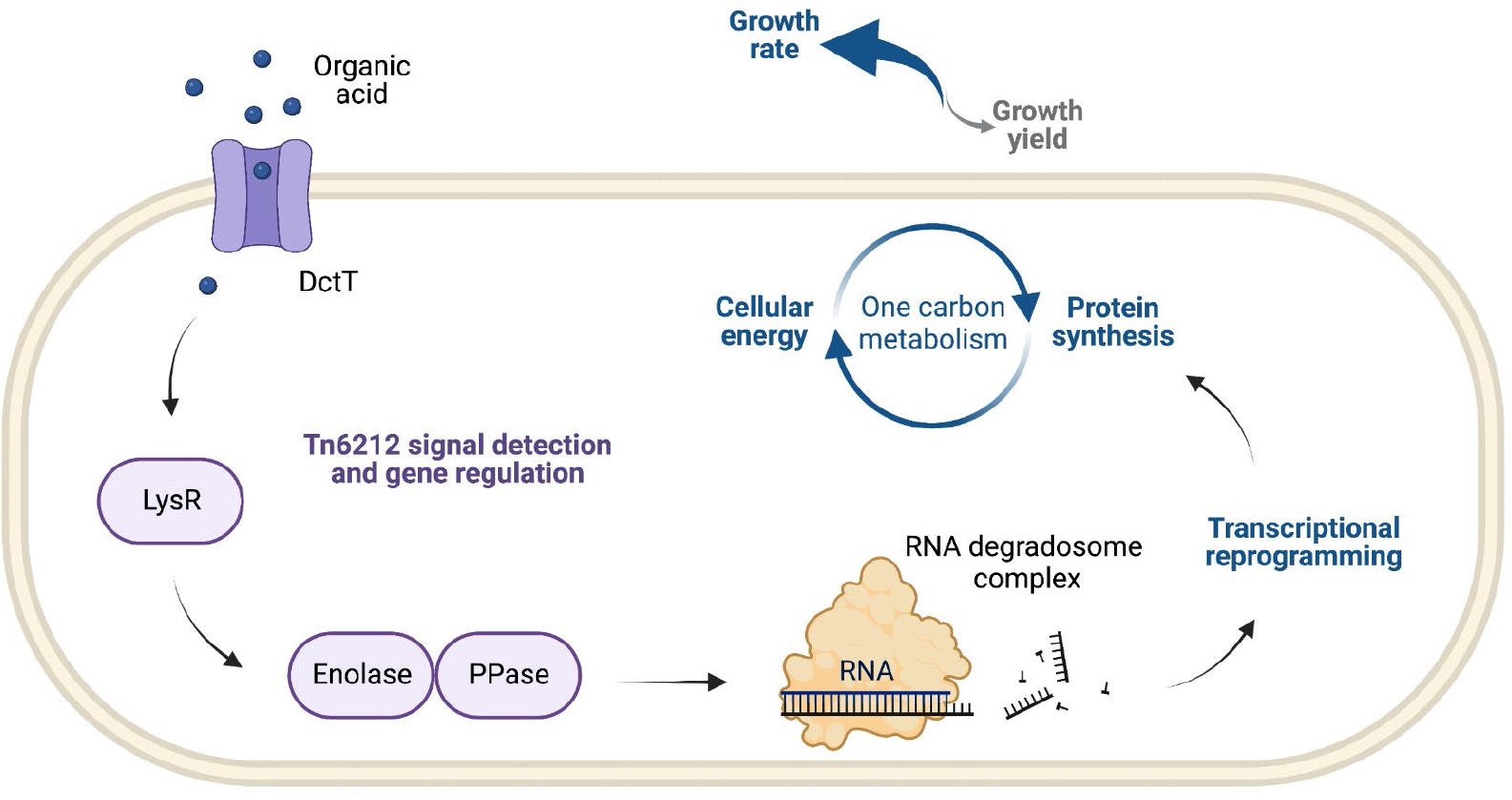
Model of Tn*6212*-ICE-bacterium-plant interactions The Tn*6212* C4-transporter DctT and transcriptional regulator LysR may sense shifts in carbon source availability and initiate a signaling cascade altering the expression of chromosomal and Tn*6212*-encoded genes. The expression of genes involved in the RNA degradosome complex, such as the genes encoding for the enolase, inorganic pyrophosphatase, RNAse E and two DEAD/DEAH box helicases, is induced. Subsequent to post transcriptional control activity of the RNA degradosome, the expression of genes involved in TCA cycle and gluconeogenesis is stimulated. This transcriptional reprogramming rapidly redistributes incoming carbon to maximize ATP, promoting growth.

The rapid redirection of cellular resources to maximize growth may enhance the fitness of bacterial strains carrying ICEs with Tn*6212*. It is possible this growth advantage also underlies ICE-mediated transposon dissemination. In a mixed population of bacteria that are either ICE- less or carrying ICEs without Tn*6212*, the subpopulation of cells carrying ICE with Tn*6212* may be overrepresented once the population reaches stationary phase, when ICE transfer is more likely to occur. Enhanced ICE dissemination - and for that matter, enhanced virulence - may be a by-product of the effect of Tn*6212* on bacterial growth in plant tissues. ICEs may therefore contribute to bacterial adaptation by adjusting bacterial transcriptional responses to preferred carbon sources.

## Acknowledgments and Funding

The authors gratefully acknowledge Zespri International Limited and Te Puke Fruit Growers Association for financial support to EC, and the Max Planck Society for generous core support to HCM and PBR. The authors thank Matt Templeton for the pMT vectors. The sponsors had no role in the design, collation, or interpretation of data.

## Supplementary Material

### Supplementary Figures

Fig. S1. Phylogeny of the strains harbouring the PsICEs.

Fig. S2. PsICEs are members of a broad family of ICEs circulating in the genus *Pseudomonas*.

Fig. S3. The backbone genes of the PsICEs.

Fig. S4. Neighbor-net network tree of backbone genes of PsICEs. Fig. S5. Conservation of Tn*6212*.

Fig. S6. Tn*6212* DctT secretion assays.

Fig S7. Growth of *Psa* NZ13 and *Psa* NZ13 ΔTn*6212* on kiwifruit. Fig. S8. Tn*6212* mutant fitness on multiple carbon sources.

Fig. S9. Correlation between genes exhibiting significant expression fold change in *Psa*

NZ13ΔTn*6212* and *Psa* NZ13 Δ*lysR*.

Fig. S10. Colony expansion rate on multiple carbon sources.

### Supplementary Tables

https://docs.google.com/spreadsheets/d/1KVfyR0oCbhGPfPfpIDWkKcjUTETS-b1bWs6emPlFJoQ/edit?usp=sharing

---

**Fig. S1.**
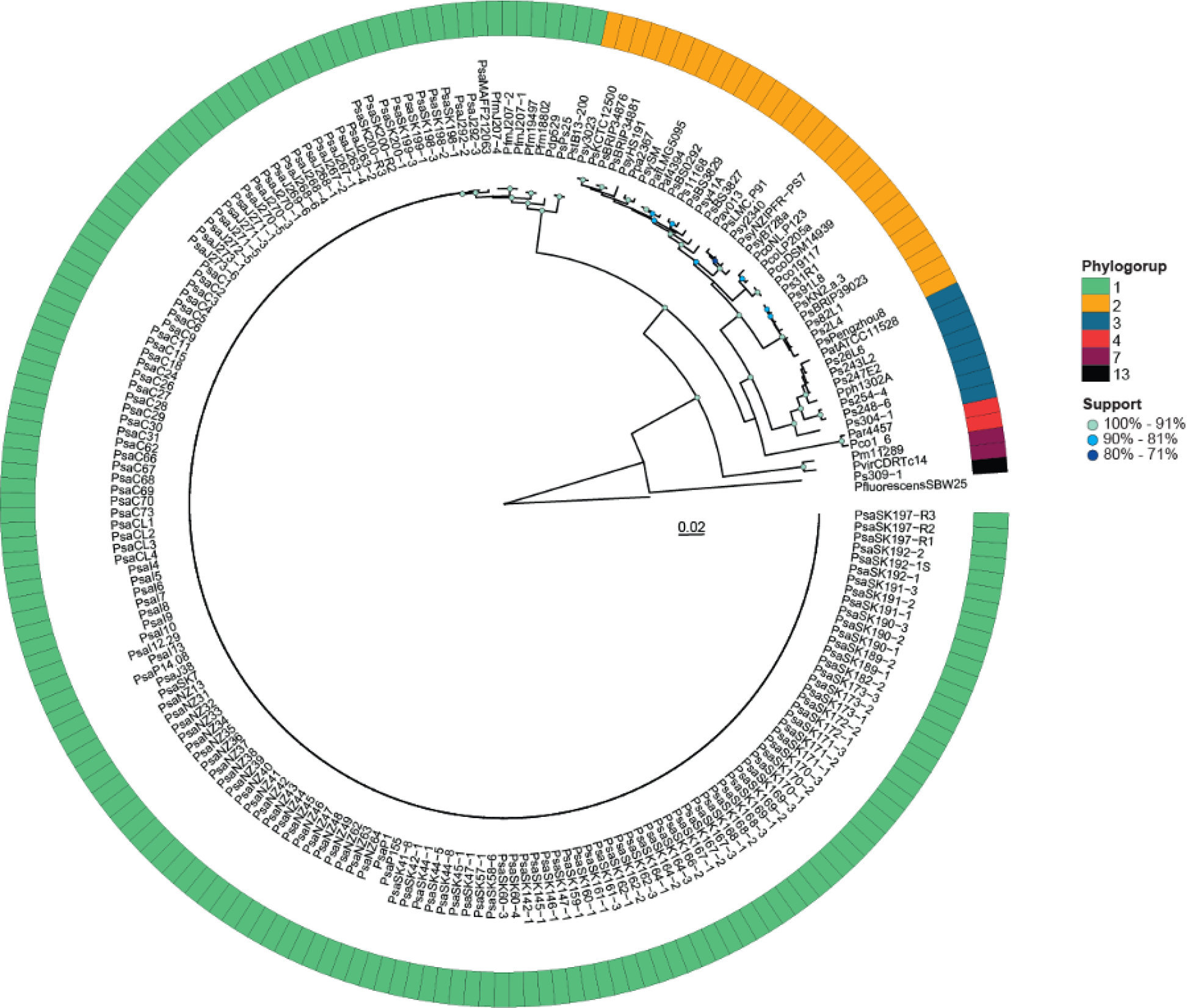
Phylogeny of the strains harbouring the PsICEs. PhyML tree based on the concatenation of *gapA*, *gltA*, *gyrB* and *rpoD*. *P. fluorescens* SBW25 was used as outgroup. The ring display the phylogroup of the *P. syringae* species complex strain. Scale bar indicates substitution per site.

**Fig. S2.**
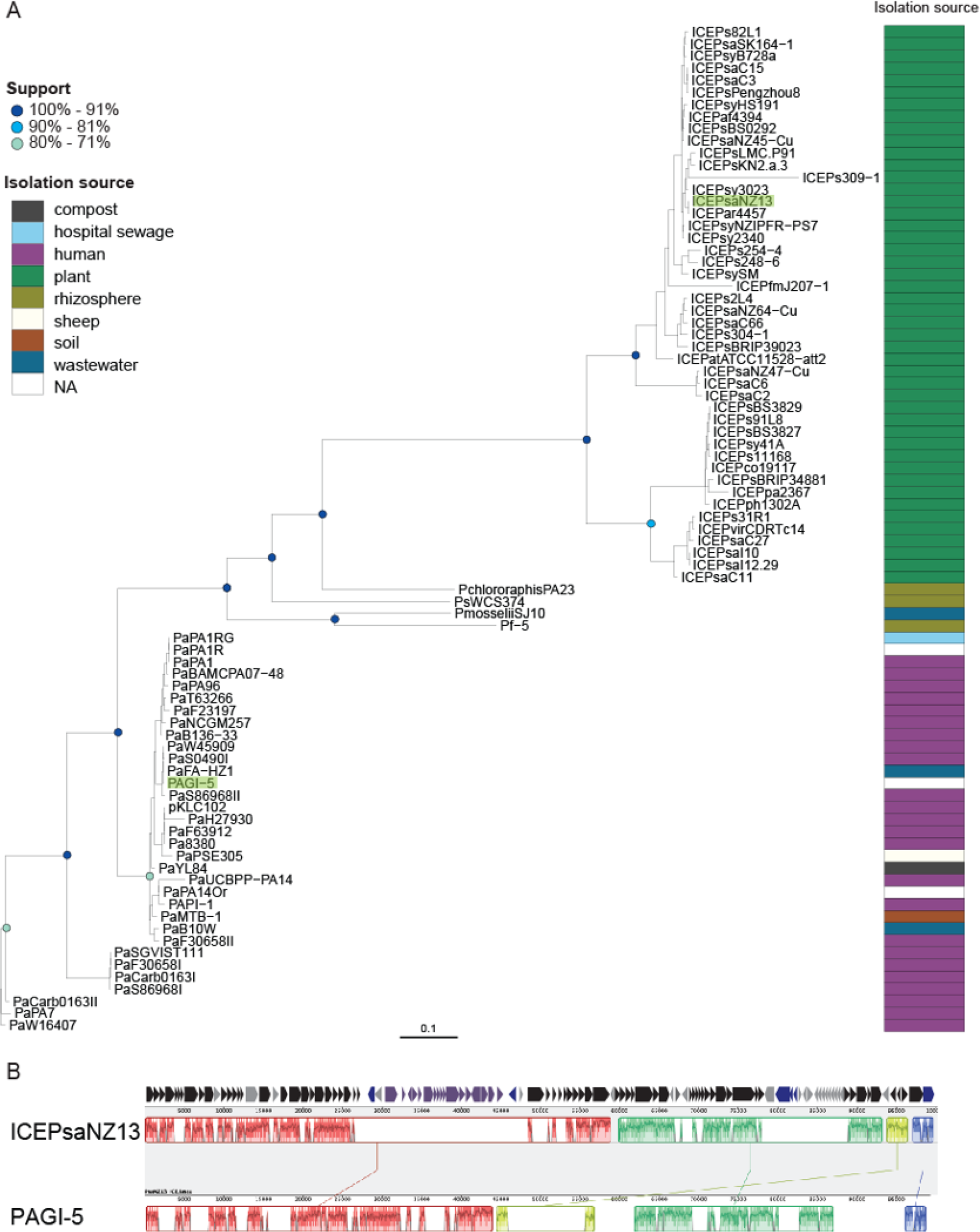
PsICEs are members of a broad family of ICEs circulating in the genus *Pseudomonas* A) PhyML tree built based on amino acid alignment of ICE proteins ParA, Topo3 and XerC. ICEs in P. syringae are named ICEStrainID, where Strain ID includes either the species (‘Ps’) or pathovar prefix (e.g. ‘Pph’), followed by the strain ID suffix (e.g. ‘1302A’). ICEs in other species are identified according to accepted names (e.g. ‘PAGI-5’, ‘pKLC102’) or using the species/pathovar prefix and strain ID suffix (e.g. ‘PaPA1’). Scale bar indicates substitution per site. B) A progressiveMauve alignment of entire ICEPsaNZ13 (representative of P. syringae ICEs) and entire PAGI-5 ICE (representative of the P. aeruginosa PAPI-1 family of ICEs).

**Fig. S3.**
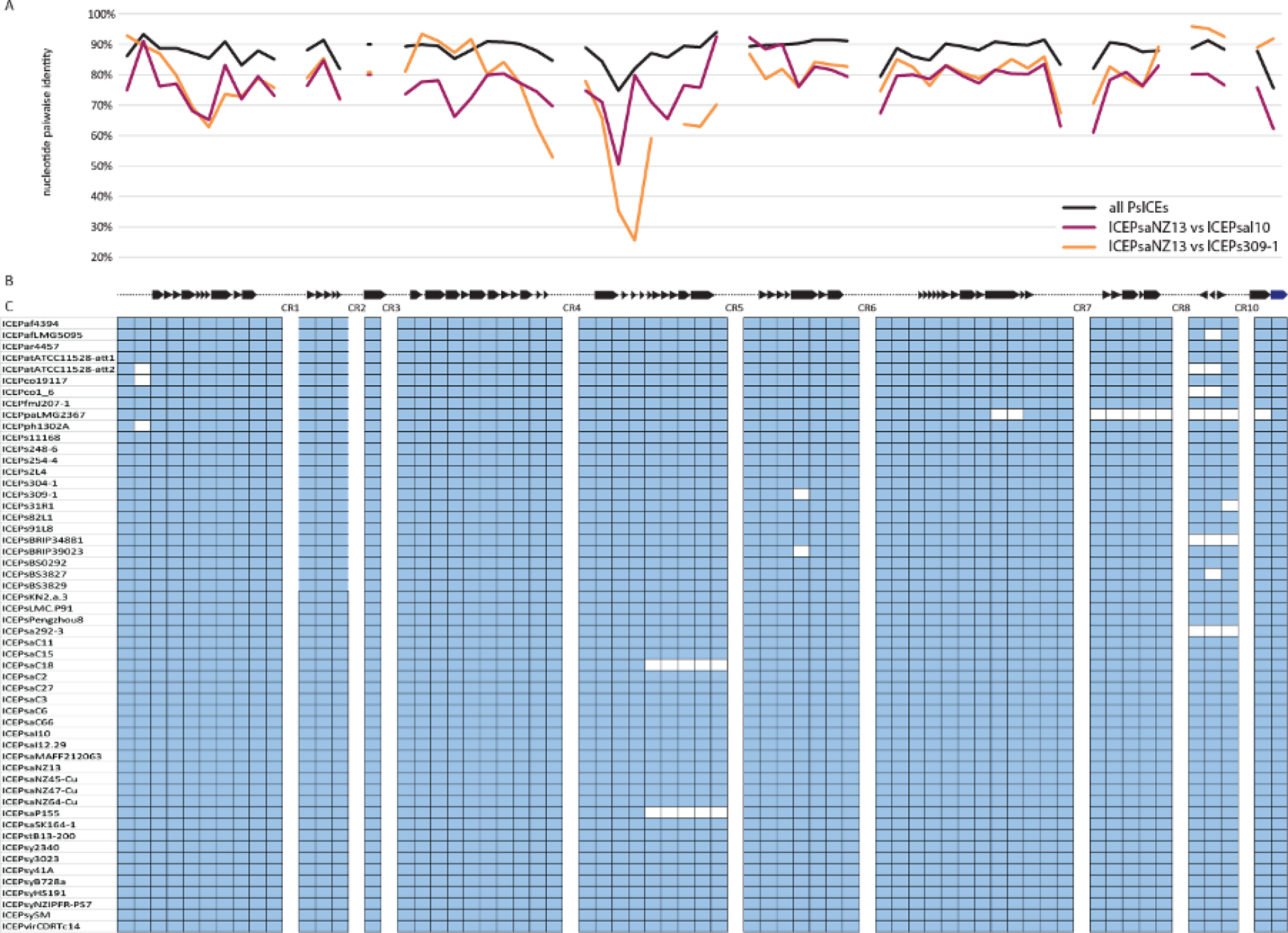
The backbone genes of the PsICEs. A) Nucleotide pairwise identity of each backbone gene. B) Graphical representation of the backbone genes. C) Presence and absence of backbone genes in the non-redundant PsICE. In light blue gene presence, in white gene absence.

**Fig. S4.**
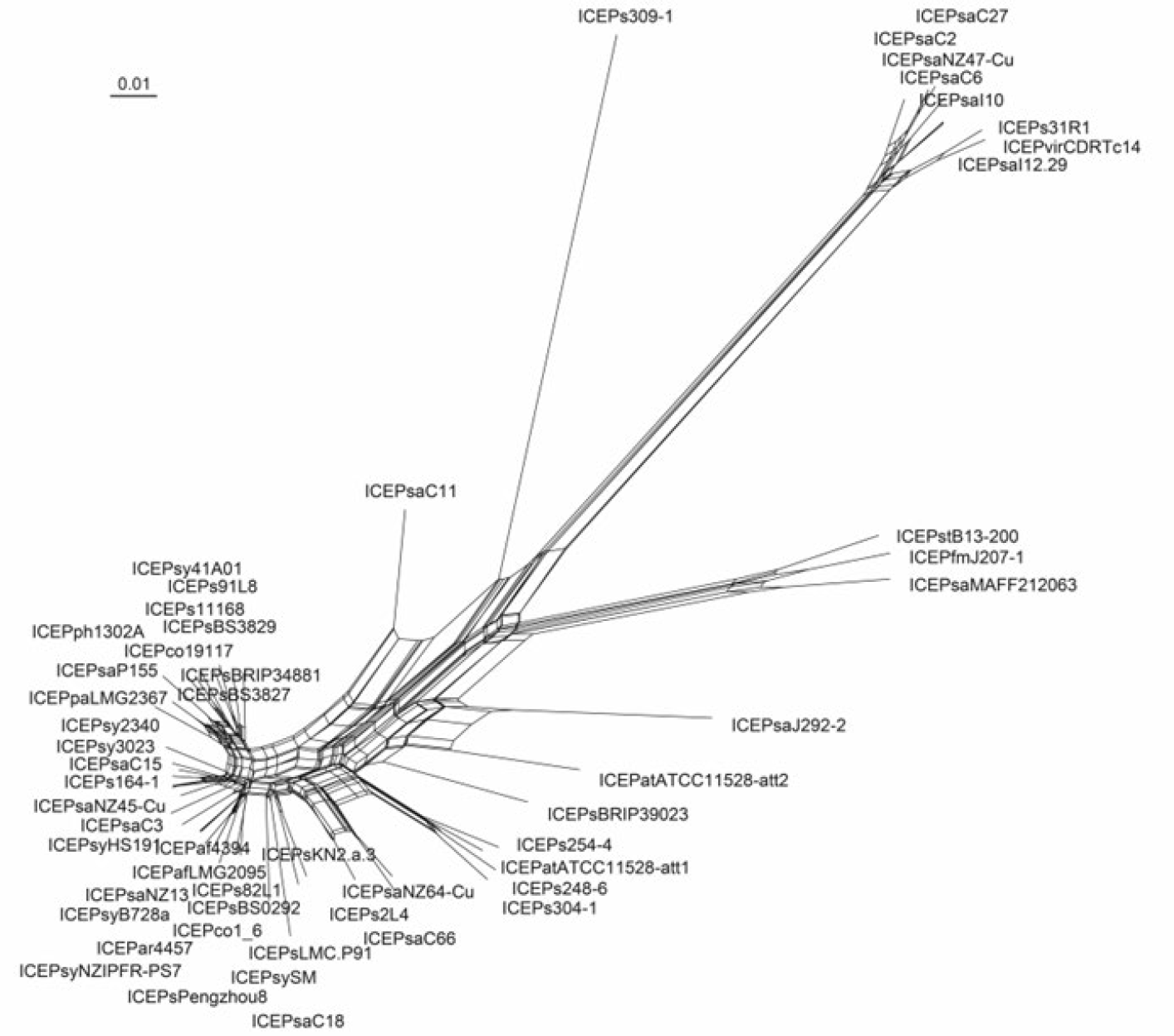
Neighbor-net network tree of backbone genes of PsICEs. Neighbor-Net generated in Splitstree using a concatenated alignment of backbone genes conserved in 53 non-redundant ICEs. Scale bar indicates substitution per site.

**Fig. S5.**
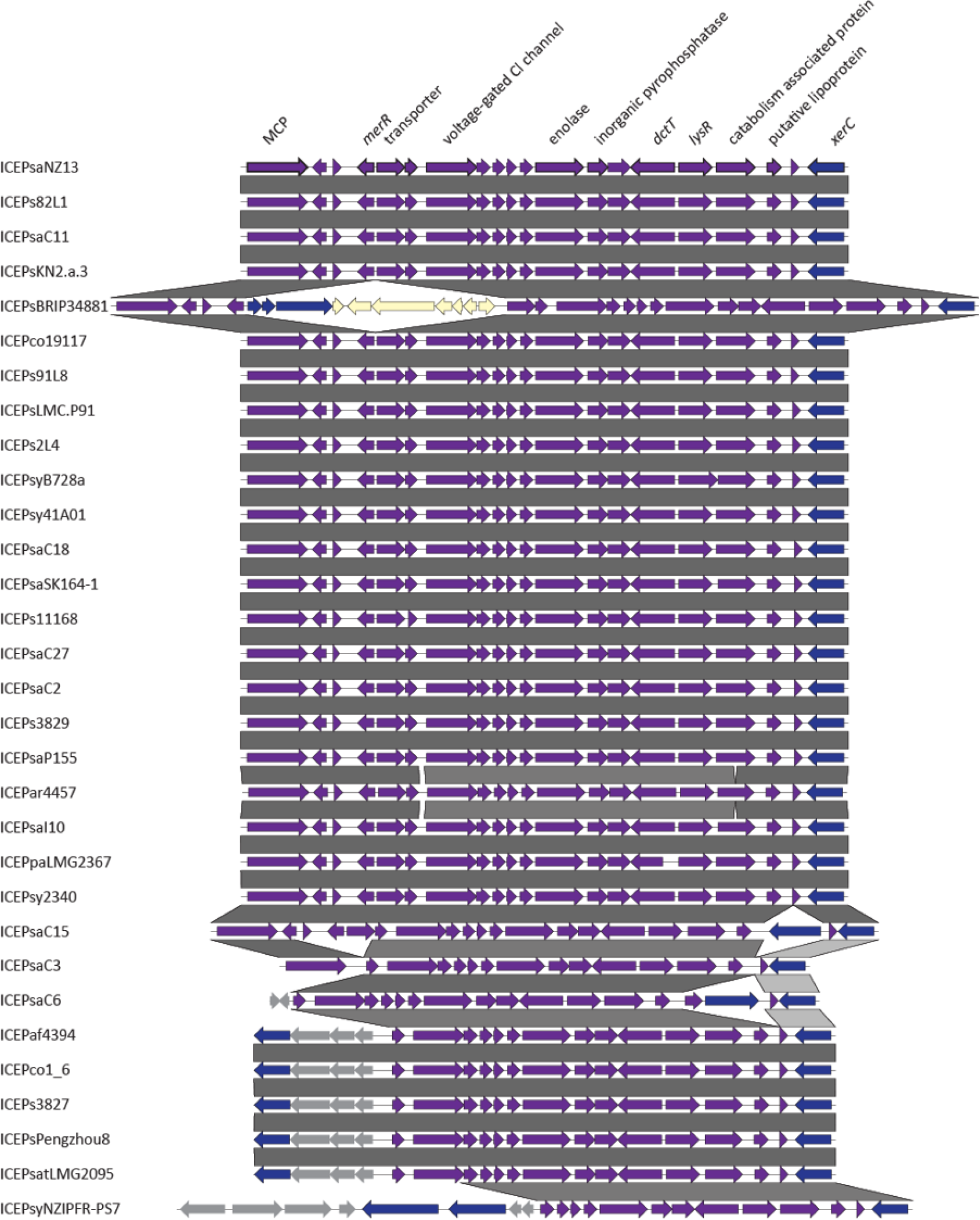
Conservation of Tn*6212*. Comparison of Tn*6212* sequences identified among the non-redundant PsICEs, generated with Easyfig v. 2.2.4. Genes are coloured as in Figure 1. Grey areas between linear structures indicate the level of their sequence identity is above 90%.

**Fig. S6.**
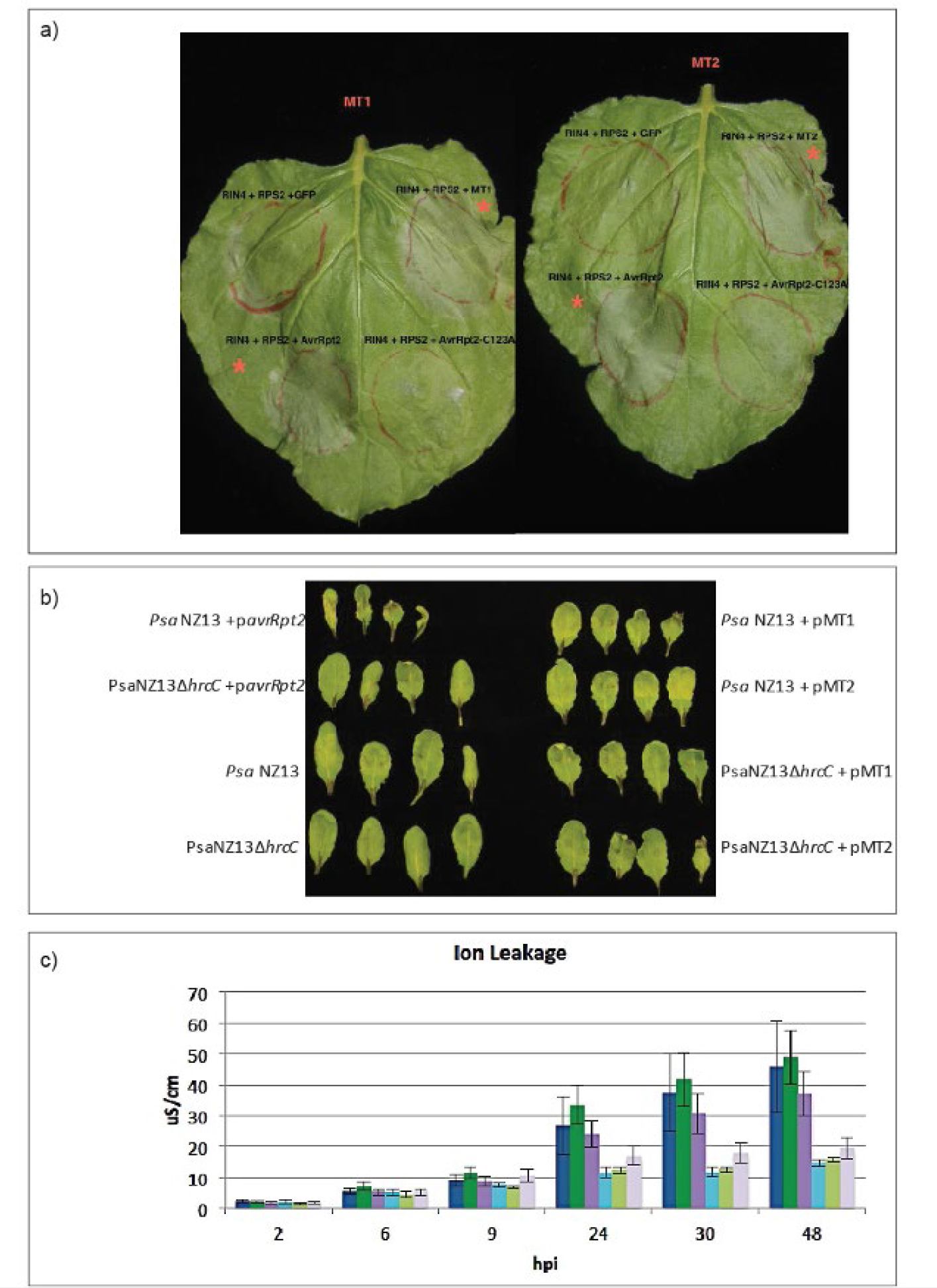
Tn*6212* DctT secretion assays. a) Agroinfiltration in *Nicotiana bethamiana* leaves using the indicated expression constructs. Pictures were taken at 24 hpi, an asterisk indicates hypersensitive response (HR) observed. b) Secretion assay in *Arabidopsis thaliana*. Leaves of *A. thaliana* Col-0 were pressure infiltrated with the indicated strain and HR development recorded at 48 hpi. The experiment was repeated two times inoculating two leaves of three plants per strain used. C) Ion leakage in *A. thaliana.* Conductivity (µS/cm) of solution containing leaf discs inoculated with *Psa* N13 (blue bars), *Psa* NZ13 + pMT-1 (dark green bars), *Psa* NZ13 + pMT-2 (purple bars), *Psa* NZ13 Δ*hrcC* (azure bars), *Psa* NZ13 Δ*hrcC* + pMT-1 (light green bars) and *Psa* NZ13 Δ*hrcC* + pMT-2 (lilac bars). Data are means and standard deviation of four replicates.

**Fig S7.**
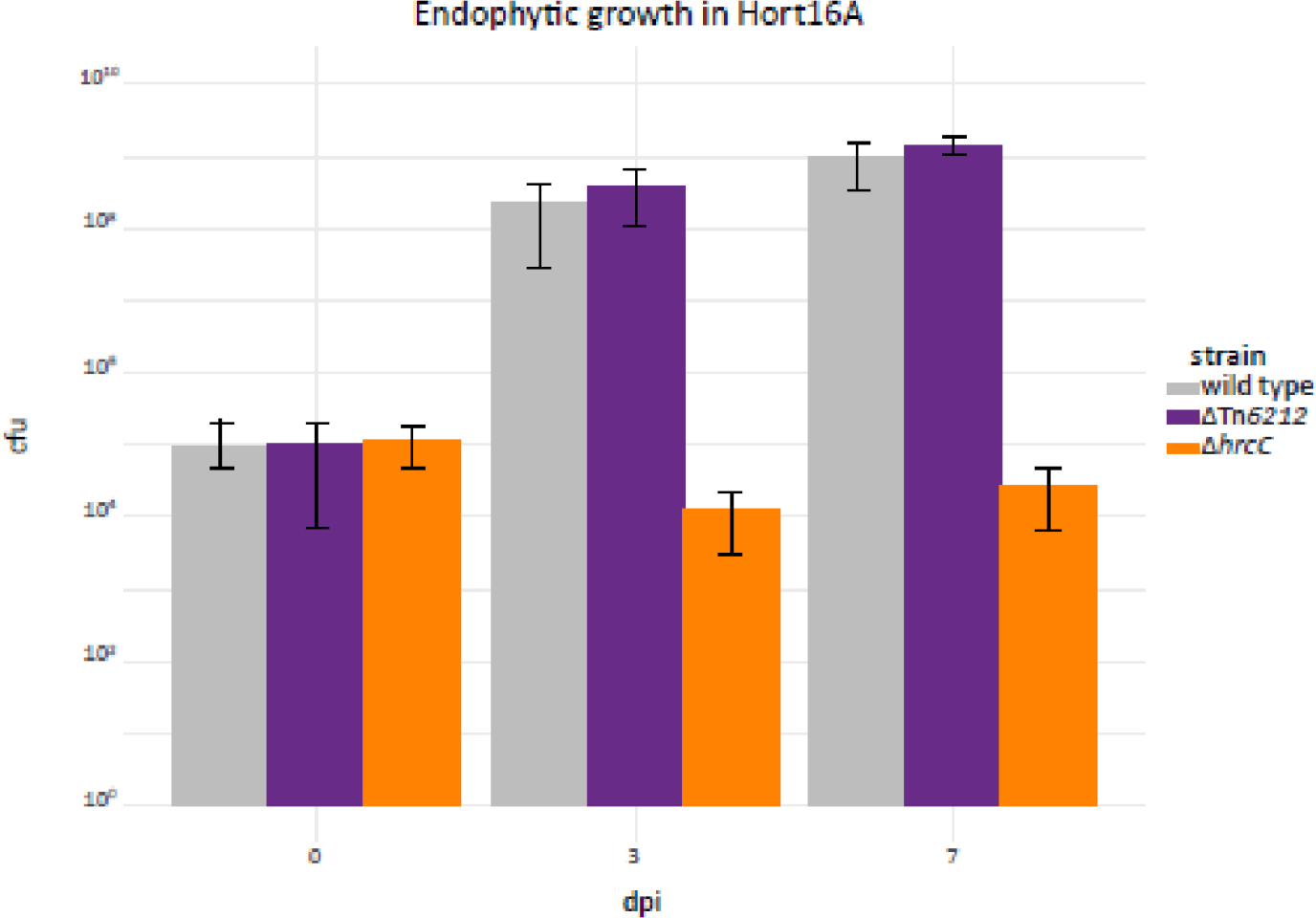
Growth of *Psa* NZ13 and *Psa* NZ13 ΔTn*6212* on kiwifruit. Growth of *Psa* NZ13 (grey bars), *Psa* NZ13 ΔTn*6212* (purple bars) and *Psa* NZ13 Δ*hrcC* (orange bars) was assessed endophytically on leaves of the kiwifruit cultivar Hort16A. Data are means and standard deviation of five replicates. Two tailed t-test revealed no statistical difference (P>0.05) between Psa NZ13 and the Tn*6212* mutants.

**Fig. S8.**
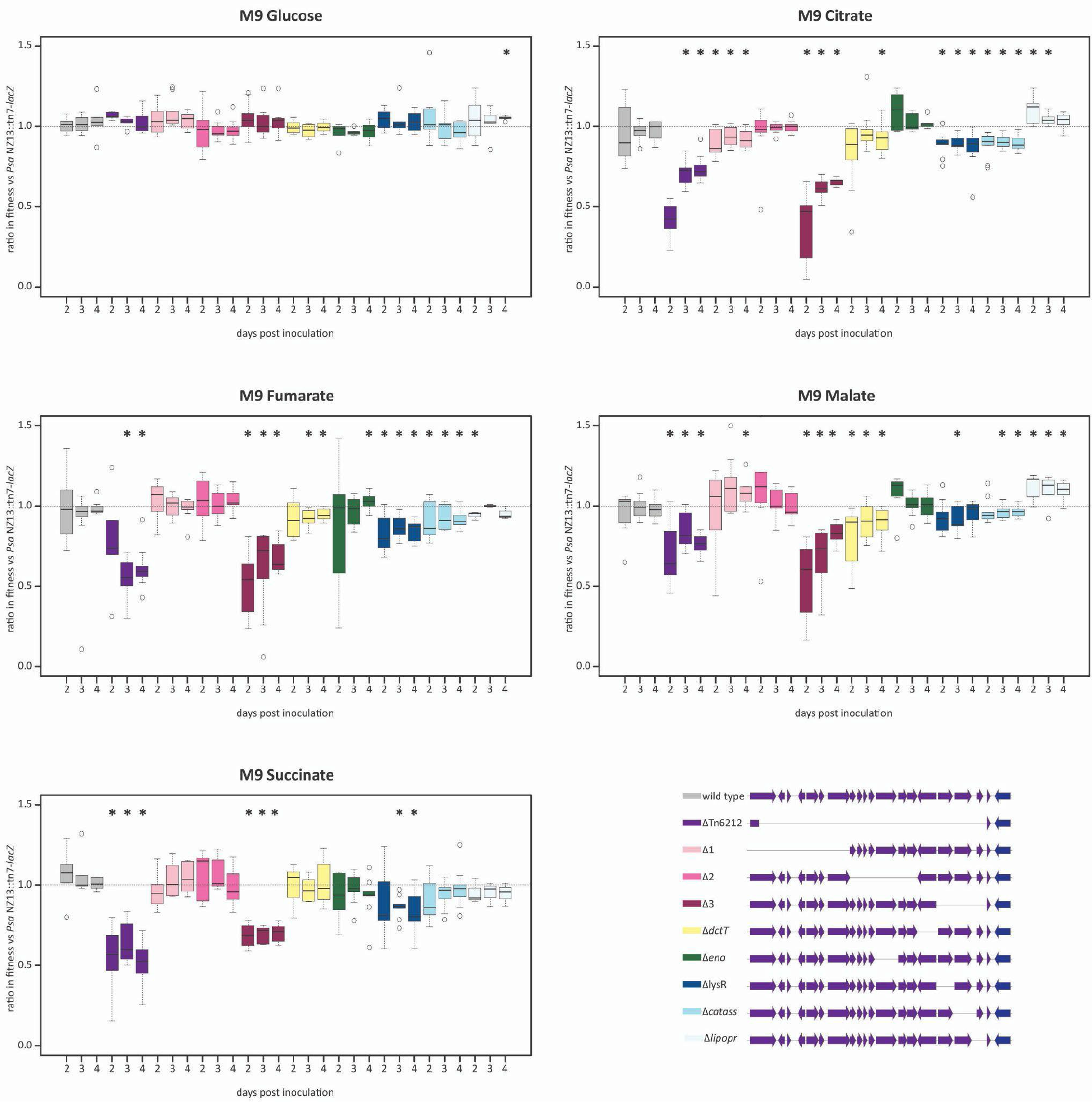
Tn*6212* mutant fitness on multiple carbon sources. Box plots of the fitness of the *Psa* NZ13 and the mutants on Tn*6212* versus *Psa* NZ::tn7-*lacZ*. Competition assays were established using equal starting densities (1:1) in M9 minimal medium supplemented with different carbon sources, measuring wildtype to mutant growth at 2, 3 and 4 days post inoculation. Values smaller than 1 indicate the competitor exhibits lower fitness relative to the wildtype strain. The experiment was performed with three replicates and repeated three times. From left to right, *Psa* NZ13 wild type, *Psa* NZ13ΔTn*6212*, *Psa* NZ13 Tn*6212*Δ1, *Psa* NZ13 Tn*6212*Δ2, *Psa* NZ13 Tn*6212*Δ3, *Psa* NZ13 Δ*dctT*, *Psa* NZ13 Δeno, *Psa* NZ13 Δ*lysR*, *Psa* NZ13 Δ*cta*, *Psa* NZ13 *Δlpr*. An asterisk indicates the fitness difference is statistically significant (one-sided one sample t-test p < 0.05).

**Fig. S9.**
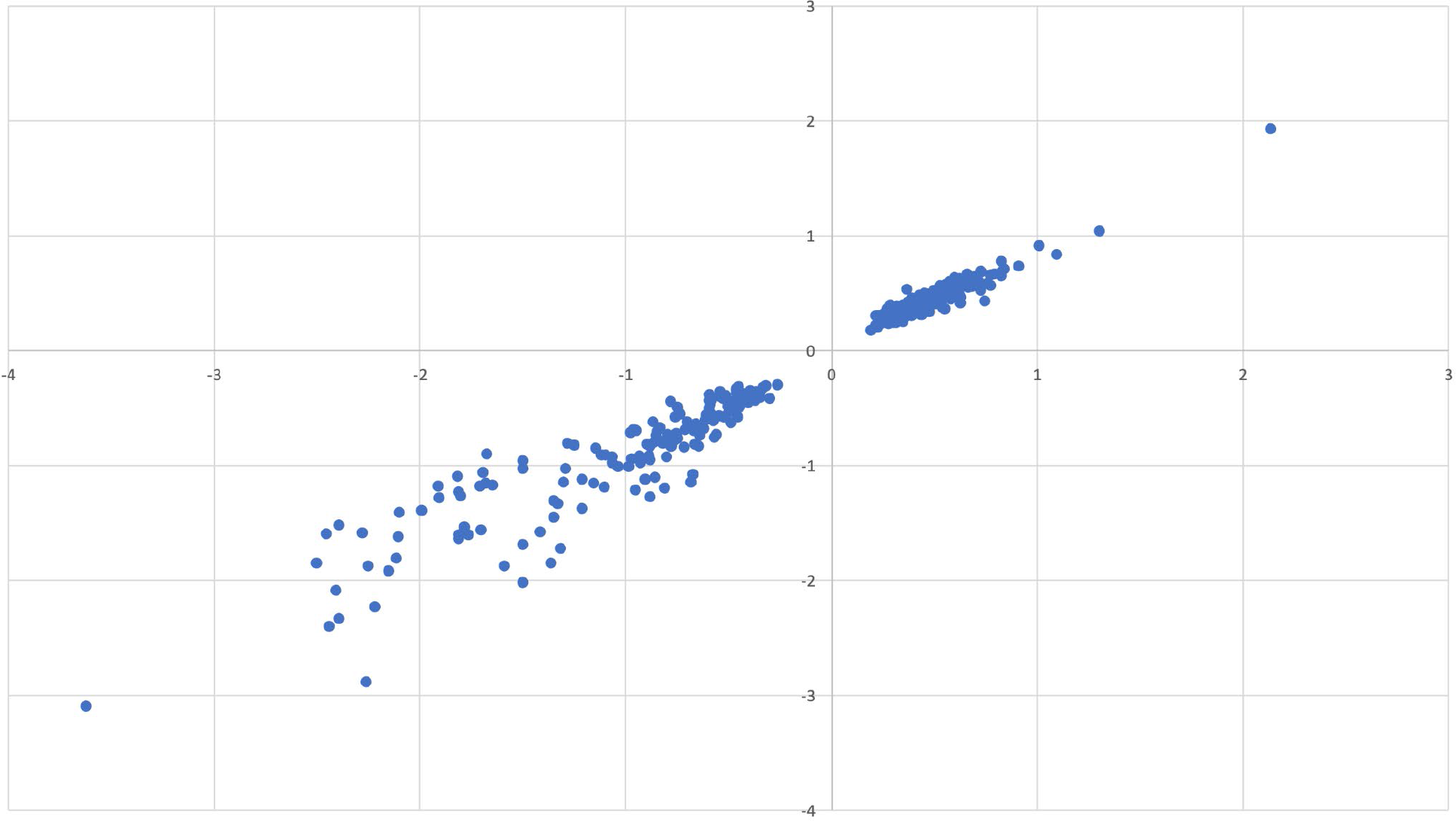
Correlation between genes exhibiting significant expression fold change in *Psa* NZ13ΔTn*6212* and *Psa* NZ13 Δ*lysR* Log2fold expression change of genes exhibiting significant differences in both ΔTn*6212* (x-axis) and ΔlysR (y-axis) strains during growth on succinate.

**Fig. S10.**
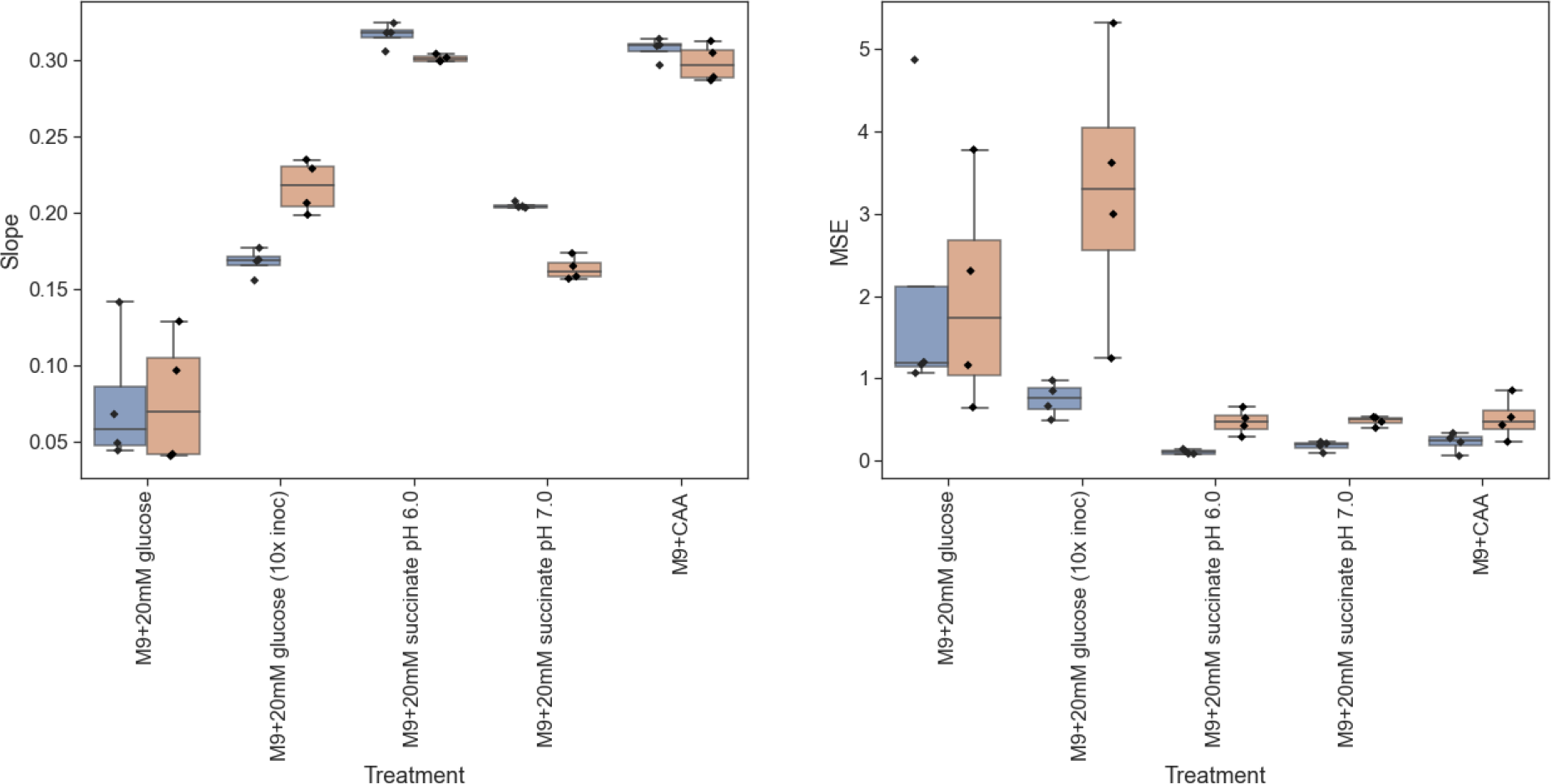
Colony expansion rate on multiple carbon sources. The dynamics of colony growth was fitted with a line (Swarm = Slope * t + Intercept). Here, we show the statistics of Slope parameter (left panel) - this is the expansion rate of colonies. The mean square error is the average squared difference between the predicted by fit and the actual data value, it is an estimator fit quality. On panels, each colony is represented by a dot. They are grouped by the treatment (x-axis) and genotype (bar color: blue - Psa, orange - ΔTn6212). A boxplot display dataset based on the five-number summary: the minimum, the maximum (shown as whiskers), the sample median (central line), and the first and third quartiles. The parameter R^2 is the fraction of the variation in data explained by the linear model. The R^2 > 0.8 for all replicates and genotypes that indicate linear relationship of the colony expansion dynamic. Statistical significance was calculated with the t-test, experiment was performed with 5 replicates.

The Statistical analysis using Student’s t-test tested whether the mean expansion rate statistically differs between genotypes for the same treatment, where H0: means of expansion rates of *Psa* and ΔTn*6212* for given condition equal and H1: means of expansion rates of *Psa* and ΔTn*6212* for given condition not equal.

| Treatment | p value |
| --- | --- |
| M9+20mM glucose | 0.968302 |
| M9+20mM glucose (10x inoc)* | 0.002232 |
| M9+20mM succinate pH 6.0* | 0.009191 |
| M9+20mM succinate pH 7.0* | 0.000043 |
| M9+CAA | 0.251976 |

The statistical analysis using a one-way ANOVA tested whether the expansion rate (Slope) for a particular genotype statistically differs between the treatments.

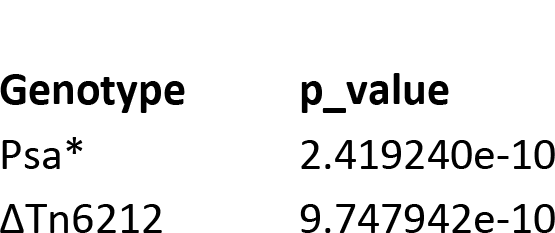

